# Metabolic Analysis of Human Retinal Pigment Epithelium and Choroid Tissue in Aging and Macular Degeneration

**DOI:** 10.64898/2026.03.24.713982

**Authors:** Emma M. Navratil, XiuYing Liu, Luke A. Wiley, Michael G. Anderson, Kacie J Meyer, Reid F. Brown, Idil A. Evan, Eric B. Taylor, Edwin M. Stone, Budd A. Tucker, Robert F. Mullins

**Author notes:** Send correspondence to: Robert F Mullins, PhD, Institute for Vision Research, The University of Iowa, 375 Newton Rd, Iowa City, IA 52242.

## Abstract

Age-related macular degeneration is a common ocular disease that causes vision loss in the elderly, with a complex set of risk factors and proposed mechanisms of pathogenesis. A powerful method for investigating changes in disease is metabolomics, by which small molecules can be identified and quantified simultaneously. We report here the metabolic analysis of human RPE-choroid tissue in aging and macular degeneration (AMD), as well as comparisons of human macular and extramacular RPE-choroid and neural retina. Levels of 215 metabolites were determined in young donors, AMD donors (early/intermediate, geographic atrophy, and neovascularization) and age-matched controls. The largest number of metabolite differences were observed between young and healthy aged controls, as opposed to between aged controls and any stage of AMD. Two notable metabolites found to be increased in aging choroids are trimethylamine N-oxide and uric acid, both of which were significant after Bonferroni correction. A mouse endothelial cell line treated with a high concentration of uric acid exhibited reduced migration in a wound closure assay. This study provides initial insights into the metabolome of human choroids in varying states of age and macular degeneration, as well as functional implications of these changes in the aging choroid.

## INTRODUCTION

Age-related macular degeneration (AMD) is a leading cause of vision loss in the elderly in industrialized countries. It is estimated that nearly 200 million people worldwide are living with some form of AMD, with that number expected to increase as the population ages (1). AMD exists in early, intermediate, and late stages, with late stage AMD divided into wet and dry AMD. In wet AMD, or macular neovascularization, blood vessels can grow into the subretinal space, resulting in tissue damage and vision loss. In dry late AMD, or geographic atrophy, large patches of the retina, retinal pigment epithelium (RPE), and choroid die causing irreversible vision loss. AMD is a complex disease with both genetic and environmental risk factors. Genetic risk factors include variants in CFH (complement factor H) (2–5) and ARMS2/HTRA1 (6–9). Major environmental risk factors include age and smoking (10–13). Options for prevention and treatment are limited to a daily vitamin regimen to modestly decrease risk of progression from intermediate to late AMD (14,15), and intraocular injections of anti-VEGF drugs to treat neovascularization (16). Recently, two complement inhibitor drugs were approved by the FDA for slowing the growth rate of geographic atrophy, though they result in increased incidence of neovascularization and do not restore vision that is already lost (17,18).

One relatively new technology being used to study many diseases, especially those with complex etiology, is metabolomics, by which hundreds of small molecules can be chromatographically identified and quantified simultaneously (19). Given that AMD is a complex disease with many risk factors and that features of the disease include the accumulation of metabolic byproducts including lipid moieties and advanced glycation end products, global metabolic study of AMD is warranted. Most previous studies of the metabolomics of AMD have utilized plasma or serum (20–27). Urine and aqueous humor samples have also been used (28,29). Several studies have found altered carnitine metabolism in AMD patients compared to controls (21,24,25,29). Other key pathways implicated include glutamine (20) and lipid metabolism (23). Though the studies listed have provided valuable and promising insight into metabolism changes during AMD, it is not known how well the differences observed systemically functionally relate to the state of the eye. In this study we utilized an extensive collection of human donor eye tissue archived at the University of Iowa Institute for Vision Research to assemble a large cohort of donors across ages and disease states for the analysis of metabolites in the choroid and RPE. Aging changes and the effects of postmortem processing time in the choroid and retina of mice were also examined. Finally, in wound closure assays, uric acid was found to inhibit endothelial cell migration.

## METHODS

### Mouse Eye Processing and Preservation

All animal experiments were conducted under a protocol approved by the University of Iowa Office of Animal Resources and were performed in accordance with the ARVO Statement for the Use of Animals in Ophthalmic and Vision Research.

Processing and preservation varied slightly between different experiments. For experiments to determine the effect of postmortem time on metabolite abundance in the eye (Sup. Fig. 1), six F1 progeny (n=3 male, n=3 female) of male *Cdh5*-cre mice (The Jackson Laboratory strain #006137) and female floxed Td-Tomato mice (The Jackson Laboratory strain #007914) at 8-weeks of age were euthanized with CO_2_ administered at 3 L/min. After confirmation of death both eyes were removed from the orbit with curved forceps. One eye was dissected immediately, with the anterior segment and lens removed and the RPE/choroid/sclera and retina separated, then flash frozen in liquid nitrogen. The second eye was placed in an Eppendorf tube and stored at 4°C for 7 hours, then dissected and frozen in the same manner as the first eye. Tissues were stored at -70°C until used in subsequent experiments.

**Figure 1.**
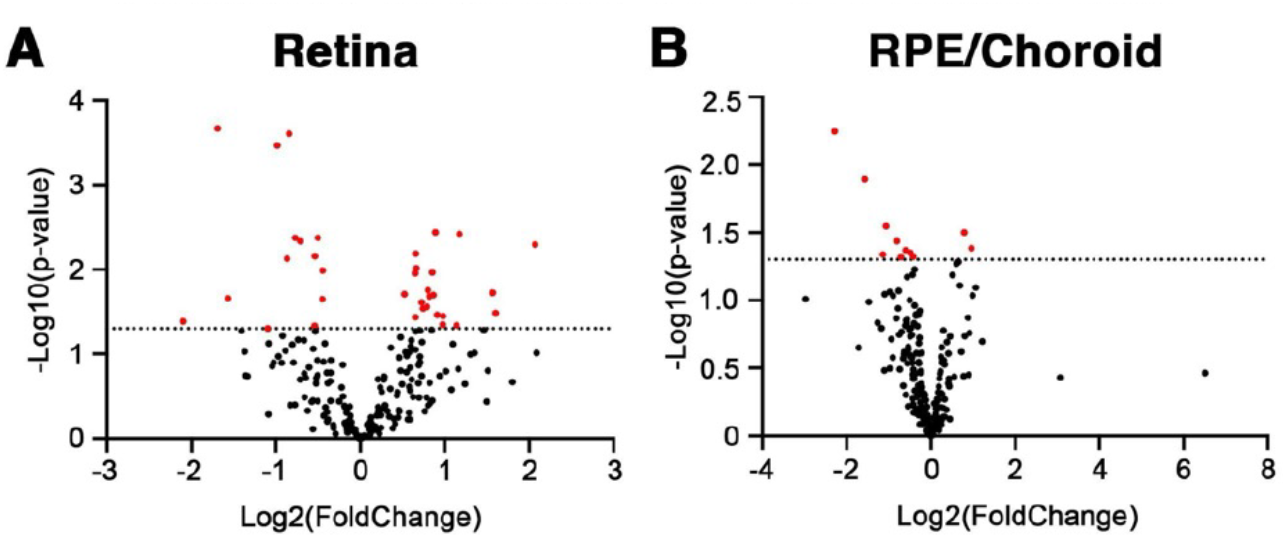
Metabolic differences in macular and extramacular human RPE/choroid and neural retina from the same donors. A volcano plot representing metabolite differences in the retina between the two regions is shown in A, and differences in the RPE/Choroid are shown in B. Dotted lines indicate a p-value of 0.05.

For assessing the effects of aging, 16 C57BL/6J mice were acquired from The Jackson Laboratory (Strain #000664) and acclimated for 9 days before the experiment (n=4 each of 8w females, 8w males, 91w females, 90w males). Mice were euthanized one at a time, out of view of cage mates, by cervical dislocation and each eye was dissected immediately with retina and RPE/choroid/sclera samples separated and flash frozen in liquid nitrogen. The time between cervical dislocation and flash freezing was 5 minutes or less.

### Human Donor Eye Processing and Preservation

Only tissue samples from deceased individuals were used in this study. Donor eyes were acquired via the Iowa Lions Eye Bank with full consent of the next of kin and in compliance with the Declaration of Helsinki. Collection of donor eyes used for this study occurred between 2005 and 2026. Tissue was obtained and processed by the laboratory within 8 hours of death. In brief, all donor eyes were processed by removing the anterior portion of the eye and ‘flowering’ the posterior pole open. Circular punches of macular (6-12mm) and peripheral (4-8mm) tissue were taken. The neural retina and RPE/choroid were separated and placed in independent tubes, then flash frozen in liquid nitrogen. Tissue was stored long term at -80 to -70 C. In contrast to mouse experiments, human RPE-choroid was separated from the sclera.

Demographic information for individual donors of tissue utilized in these experiments can be found in Supplemental Table 1, while summary demographic data can be found in Table 1. Death-to-preservation (freezing) time did not differ between study groups.

**Table 1.** Demographics Summary Table.

| **Group** | **Male** | **Female** | **Age Range** | **Age (mean, median)** | **DtoP Time (mean, median)** | **Total n** |
| --- | --- | --- | --- | --- | --- | --- |
| Young | 4 | 4 | 21-42 | 32.25, 34 | 6h 8m, 6h 6m | 8 |
| Aged Control | 17 | 20 | 70-97 | 83.73, 85 | 5h 48m, 5h 45m | 37 |
| AMD | 11 | 10 | 71-96 | 84.71, 85 | 5h 52m, 6h 6m | 21 |
| nAMD | 5 | 9 | 70-97 | 83.86, 83 | 5h 41m, 5h 26m | 14 |
| GA | 0 | 7 | 76-97 | 88.43, 87 | 5h 24m, 4h 56m | 7 |
| Macula vs Periphery | 5 | 5 | 13-77 | 55.1, 65 | 6h 2m, 5h 57m | 10 |
| All | 42 | 55 | 13-97 | 77.1, 83 | 5h 50m, 5h 55m | 97 |

The eyes from the older age group were categorized as being unaffected (n=37), or having early/intermediate AMD (n=21), geographic atrophy (n=7), or macular neovascularization (n=14) based on histological examination and/or chart review by a board-certified ophthalmologist. In one donor (#82), eyes were discordant with geographic atrophy and macular neovascularization in contralateral eyes.

### Metabolomics

Sample processing: Tissue samples were lyophilized and transferred to ceramic bead tubes. Extraction solvent containing 9 internal standards was added 100-fold (w/v) to each sample followed by homogenization and rotation for one hour at -20°C. After this, samples were centrifuged at 21,000xg for 10 minutes, the supernatant was transferred to new 1.5ml microcentrifuge tubes and vortexed, then 300 ul of supernatant from each sample was moved to new tubes. Supernatant samples were dried using a speed vacuum apparatus. The dried extracts were reconstituted with 30 ul 1:1 v/v acetonitrile/water, vortexed, and stored at -20C overnight. Reconstituted extracts were centrifuged and transferred to LC-MS autosampler vials for analysis. A Thermo Q Exactive hybrid quadrupole Orbitrap mass spectrometer with a Vanquish Flex or Vanquish Horizon Ultra High Performance Liquid Chromatography system. Millipore SeQuant ZIC-pHILIC LC columns (2.1 x 1150 mm, 5µm particle size) and ZIC-pHILIC guard columns (20x2.1mm) were used with an injection volume of 2 µL. The mobile phase was composed of Solvent A (20 mM ammonium carbonate and 0.1% ammonium hydroxide (v/v), pH ∼9.1) and Solvent B (acetonitrile). The method was run at 0.150 mL/min, with a gradient starting at 80% Solvent B and decreasing to 20% Solvent B over 20 minutes, returning to 80% Solvent B over 0.5 minutes, and held at 80% for 7 minutes. Prior to each sample injection there was a 2-minute equilibration time, during which the flow was increased to 0.3 mL/minute. Absolute quantification of TMAO and uric acid was conducted similarly, with the addition of a heavy carbon standard.

### Data Analysis and Statistics

Processing of raw LC-MS data was performed with Thermo Scientific TraceFinder 5.2 software and metabolites were identified based on the University of Iowa Metabolomics Core facility standard-confirmed, in-house library. Signal drift was corrected with NOREVA method (30) and data were normalized to the sum of all measured metabolite ions in each sample. For each sample group comparison, p-values for each metabolite were determined in Excell using T-tests (two-tailed, unequal variance). For the purposes of discussion, p < 0.05 was considered significant. The Benjamini-Hochberg procedure (31) was also used to control the false discovery rate (FDR) and significance according to this procedure is indicated in Tables 2-4. Graphpad Prism Version 10.4.1 was used to generate volcano plots using the log2(foldchange) and - log10(p-value) for each metabolite.

**Table 2.** Significantly different metabolites in human extramacular tissue compared to macular tissue. Data corresponds to Figure 1. Asterisks (*) indicate significance according to Benjamini-Hochberg FDR correction.

| Human Macula vs Extramacula |  |  |
| --- | --- | --- |
| Retina |  |  |
| Metabolite | P-value | Log2(FoldChange) |
| Phosphoenolpyruvic acid | 0.0002 * | -1.69 |
| N-Acetylglutamic acid | 0.0003 * | -0.84 |
| S-Adenosylhomocysteine | 0.0003 * | -0.98 |
| Deoxyadenosine monophosphate | 0.0036 | 0.89 |
| Argininosuccinic acid | 0.0038 | 1.17 |
| N-Acetylornithine | 0.0042 | -0.77 |
| Flavin adenine dinucleotide | 0.0042 | -0.50 |
| N-Acetylaspartic acid | 0.0046 | -0.71 |
| N-methyl-D-aspartate | 0.0051 | 2.07 |
| Glycine | 0.0065 | 0.66 |
| Creatine | 0.0070 | -0.54 |
| Glutamic acid | 0.0075 | -0.87 |
| Adenine deoxyribonucleoside | 0.0097 | 0.66 |
| Alanine | 0.0103 | -0.44 |
| Beta-Alanine | 0.0103 | -0.44 |
| Sarcosine | 0.0103 | -0.44 |
| Aconitic acid | 0.0108 | 0.85 |
| Dehydroascorbic acid | 0.0108 | 0.85 |
| Glutamine | 0.0111 | 0.65 |
| Kynurenine | 0.0176 | 0.80 |
| 3-Methyl-2-oxobutanoic acid | 0.0188 | 1.57 |
| Melatonin | 0.0189 | 1.56 |
| Serine | 0.0196 | 0.52 |
| Isoleucine | 0.0200 | 0.87 |
| Leucine | 0.0208 | 0.82 |
| Coenzyme A | 0.0219 | -1.57 |
| Nicotinamide adenine dinucleotide | 0.0224 | -0.45 |
| Imidazoleacetic acid | 0.0244 | 0.73 |
| Homoserine | 0.0276 | 0.79 |
| Thymidine monophosphate | 0.0289 | 0.74 |
| 4-Methyl-2-oxopentanoic acid | 0.0327 | 1.60 |
| Phenylalanine | 0.0344 | 0.91 |
| N-Acetylalanine | 0.0355 | 0.98 |
| Methionine | 0.0368 | 0.66 |
| Azelaic acid | 0.0411 | -2.10 |
| 3-Hydroxypropionic acid | 0.0448 | 0.98 |
| Lactic acid | 0.0448 | 0.98 |
| Glyceric acid | 0.0458 | 1.14 |
| Allothreonine | 0.0467 | -0.54 |
| Threonine | 0.0467 | -0.54 |
| AICAR | 0.0500 | -1.09 |
| **Choroid** |  |  |
| **Metabolite** | **P-value** | **Log2(FoldChange)** |
| N-Acetylornithine | 0.0057 | -2.28 |
| Thymidine monophosphate | 0.0128 | -1.56 |
| Uracil | 0.0284 | -1.06 |
| 3-Methylhistamine | 0.0316 | 0.79 |
| Inosine | 0.0367 | -0.81 |
| Imidazoleacetic acid | 0.0412 | 0.97 |
| CDP-Ribitol | 0.0427 | -0.59 |
| Adenine | 0.0445 | -0.49 |
| Dihydroxyacetone phosphate | 0.0458 | -1.13 |
| N-Acetylglutamic acid | 0.0477 | -0.42 |
| Adenosine monophosphate | 0.0479 | -0.71 |

Metaboanalyst 6.0 was used to generate pathway analysis for each comparison. Concentration tables were uploaded as CSV files and data was normalized by median and scaled using the auto-scaling option (mean-centered and divided by the standard deviation of each variable). The visualization method was Scatter Plot, the enrichment method was Global Test, the topology measure was Relative-betweenness Centrality, and the reference metabolome was *Homo sapiens* (KEGG).

### Wound Healing Assays

C166 immortalized mouse endothelial cells were grown in 24 well plates in DMEM high glucose with 10% heat inactivated FBS and 100 µg/mL Primocin. When cells reached confluency, scratches were made in each well using a BioTek Autoscratch wound making tool. Each well was washed once in sterile calcium/magnesium-free PBS, after which new media containing each treatment was added. Treatments included 50 µg/mL succinylated Concanavalin A lectin, 0.5mM and 1mM trimethylamine N-oxide (Millipore Sigma #317594), 100 µg/mL and 200 µg/mL uric acid, 1:500 ultrapure water (vehicle control for TMAO), and 1:250 1N NaOH (vehicle control for uric acid).

The plate was placed on a Lumascope live image microscope (Etaluma, Carlsbad, CA) housed in an incubator (37°C, 5% CO_2_) with an automated stage and a 4x Olympus objective, and a Lumascope software imaging protocol was used to image a region of each scratch once an hour. Images were analyzed using a wound healing recipe in the Lumaquant software with Low Detection (Background Removal Factor 60, Contrast Threshold 85, Fill Holes Size 10000, Smoothing Factor 10) and Medium Subset Filtering (Minimum Object Size 10000), then manually corrected for accuracy. Wound area from every 4 hours was plotted and the percentage of wound closure was determined in Excel by comparing the area at each timepoint to the area at time zero. All treatments were performed in triplicate on each plate. In addition, 3 plates of cells were run for each assay. Data were normalized as percent closure at each timepoint and were pooled for each treatment. Statistical significance was determined Excel using the T.TEST function (two-tailed and unequal variance).

## RESULTS

### Mouse: Postmortem Time Effects

An initial experiment was performed to determine the metabolic changes that occur in the retina and choroid after death, mimicking a relatively unfavorable timeframe within which the human eyes are processed (<8h) and longer than the average preservation time (Table 1). Mouse eyes were collected and dissected either immediately after CO2 euthanasia or after seven hours at 4°C. LC-MS analysis detected 236 metabolites. Of these, 110 metabolites were significantly different in the retina and 54 were significantly different in the choroid (Sup. Fig. 1), indicating more metabolic stability in the choroid than the retina during postmortem storage. The pathway analysis also reflects this, with 53 pathways identified as different in the retina after 7h versus 28 pathways identified as different in the choroid. These results are consistent with the literature regarding metabolite stability during postmortem delays before preservation. The results of this experiment provide valuable information about metabolite stability, specifically in the tissues of interest.

### Human: Macula vs Periphery

An additional preliminary experiment was conducted to compare macular and peripheral tissue from the same eye, to determine how closely the peripheral choroidal biochemistry matches that in the macula, at least in eyes without maculopathy. Macular tissue from donors 11-96 was used in a previous experiment, but peripheral tissue was still available. A cohort of 10 donors of varying ages were selected for a comparison of RPE/choroid and retinal tissue between the macula and superonasal quadrant. Metabolomic analysis identified 230 metabolites, of which 41 metabolites were significantly different in retina, while only 11 were significantly different between the two regions in the choroidal tissue (Fig. 1). Pathway analysis revealed 27 pathways different in the retina, while none were significantly different in the RPE/choroid. These results indicate that the macular and peripheral choroid are very similar in their identified metabolites, and that changes in the peripheral choroid during aging and AMD is a reasonable surrogate for the macula.

### Mouse: aging

A comparison of young and aged mouse neural retina and RPE/choroid/sclera was conducted to provide a reference point for the changes occurring in these tissues in a model that allows for low postmortem delays (<= 5mins) and low variability in genetics, environment, diet, and other factors compared to human tissue. LC-MS metabolomic analysis of neural retina and RPE/choroid/sclera in young and aged mice identified 207 metabolites, of which 24 were significantly different in the retina and 37 in the choroid (Fig. 2). Pathway analysis identified 5 pathways in the retina and 26 in the choroid that were significantly changed in the older mice compared to the younger ones. This is a reverse trend of the comparisons between macula and periphery and postmortem times, and suggests choroidal metabolism is altered to a greater degree in aging than that of the retina.

**Figure 2.**
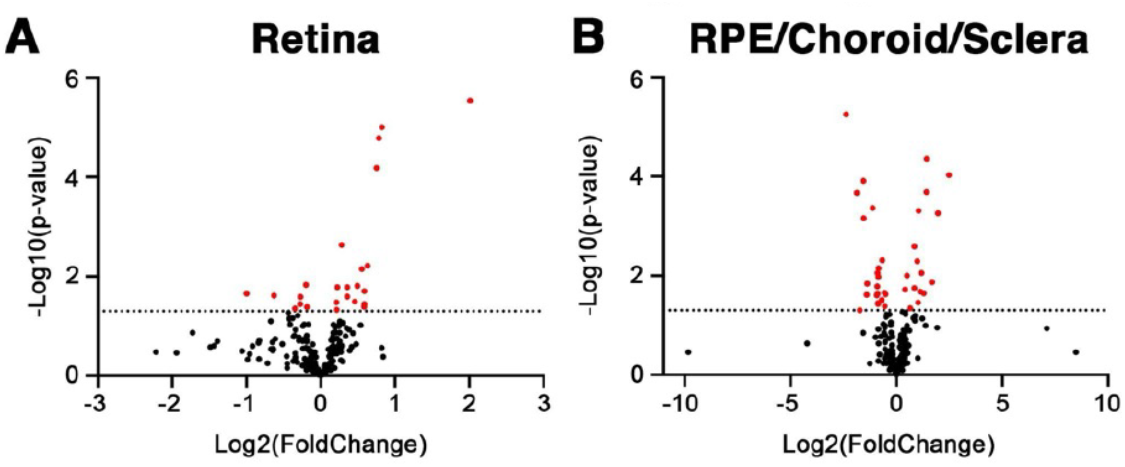
Metabolic changes in C57BL/6J mouse RPE/choroid/sclera and neural retina during aging. A volcano plot representing metabolite differences in the retina between the two age groups (8 weeks vs 90-91 weeks) is shown in A and the same comparison for RPE/choroid/sclera is shown in B. Dotted lines indicate a p-value of 0.05.

### Human: young vs aged vs AMD

Analysis of the large cohort revealed significantly more differences between young and aged RPE/choroids than between unaffected aged choroids and any stage of AMD (Fig. 3). In aged choroids, 32 metabolites and 15 pathways were significantly different from young choroids. The top five pathways were glycerophospholipid metabolism (5/36 hits, p=1.79E-03, I=0.17), mannose type O-glycan biosynthesis (1/17 hits, p=5.56E-03, I=0.06), butanoate metabolism (6/15 hits, p=6.23E-03, I=0.031), glycerolipid metabolism (3/16 hits, p=0.014, I=0.137), and purine metabolism (22/70 hits, p=0.020, I=0.47). Other significant pathways with high impact include arginine and proline metabolism, histidine metabolism, nicotinate and nicotinamide metabolism, and alanine, aspartate, and glutamate metabolism.

**Figure 3.**
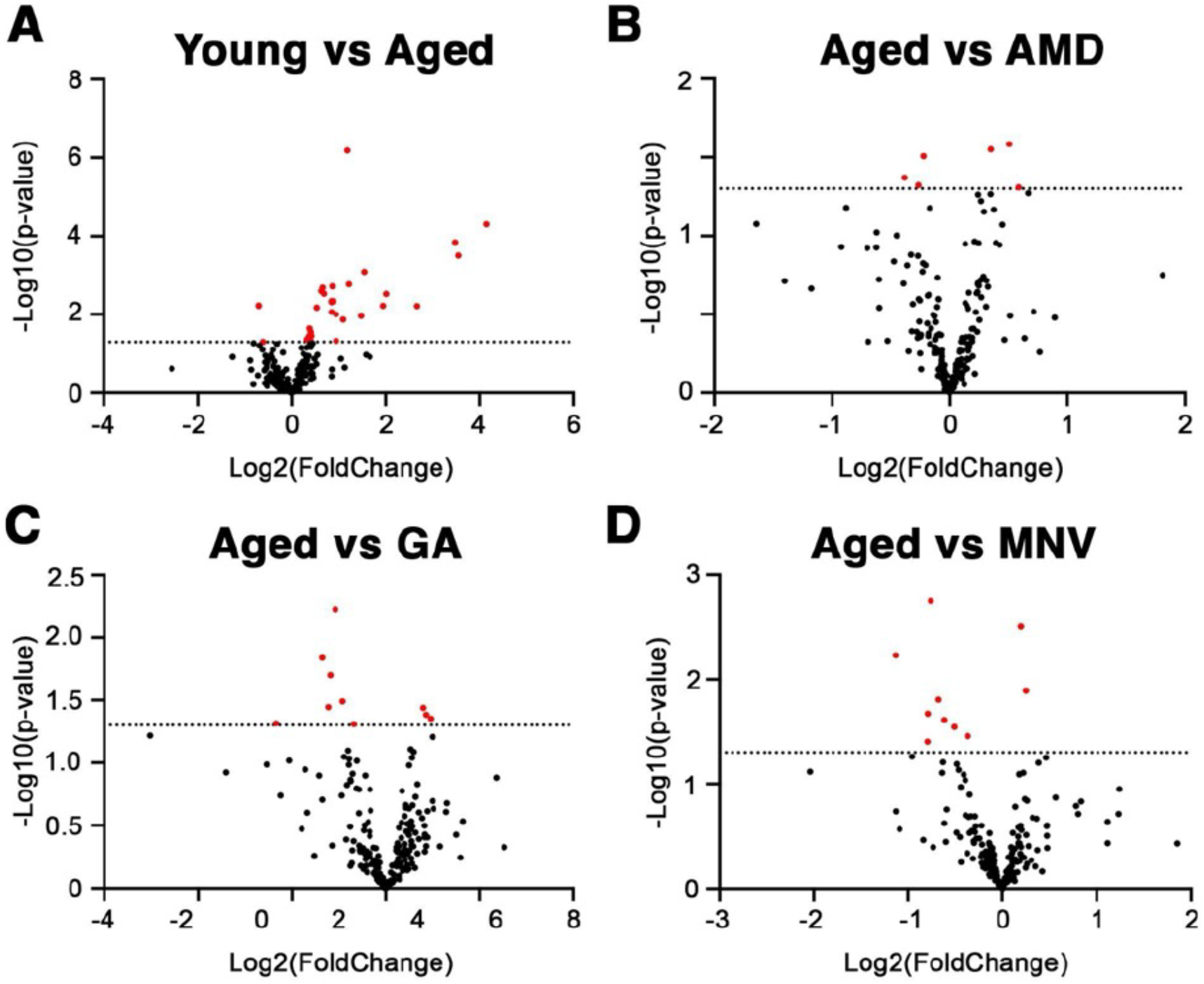
Metabolomics of human RPE/choroid in age and disease. Volcano plots of changes in metabolites are shown for young vs aged (A), aged vs early/intermediate AMD (B), aged vs geographic atrophy (C), and aged vs neovascular AMD (D). Dotted lines indicate a p-value of 0.05.

Compared to unaffected aged choroids, choroids with early/intermediate AMD (Fig. 3B, Table 3) showed differences in 6 metabolites (N-acetylneuraminic acid, glutamic acid, histidine, 3-methylhistamine, N-formylmethionine, and arachidonoyl-L-carnitine) and 3 pathways (nitrogen metabolism, porphyrin metabolism, and glycine, serine, and threonine metabolism); those with geographic atrophy (Fig. 3C) were distinct in 11 metabolites (azelaic acid, nonanoic acid, heneicosanoic acid, stearic acid, heptadecanoic acid, arachidic acid, N-acetylglutamic acid, glycine, S-adenosylhomocysteine, glutarylcarnitine, and pentadecanoic acid) but no pathways; and choroids from eyes with macular neovascularization (MNV) (Fig. 3D) displayed differences in 10 metabolites (dAMP, aminolevulinic acid, TMP, glutamine, GMP, arachidonoyl-L-carnitine, AMP, adenine deoxyribonucleoside, GDP, and CDP) and 3 pathways (steroid hormone biosynthesis, pyrimidine metabolism, and porphyrin metabolism). Notably, several of these metabolites such as azelaic acid and odd-carbon fatty acids are likely to be derived from the microbiome.

**Table 3.** Metabolites with altered abundance in aging mouse retina and RPE/choroid. Data corresponds to Figure 2. Asterisks (*) indicate significance according to Benjamini-Hochberg FDR correction.

| **C57BL/6J Young vs Aged** |  |  |
| --- | --- | --- |
| **Retina** |  |  |
| **Metabolite** | **P-value** | **Log2(FoldChange)** |
| CAR(5:0-DC) | 2.89E-06 * | 2.01 |
| Carnitine | 9.92E-06 * | 0.82 |
| Acetyl-L-carnitine | 1.66E-05 * | 0.78 |
| CAR(4:0) | 6.57E-05 * | 0.75 |
| CAR(3:0) | 0.0023 | 0.28 |
| Pantothenic acid | 0.0061 | 0.63 |
| CAR(12:0) | 0.0072 | 0.56 |
| Proline | 0.0149 | -0.20 |
| Valerylcarnitine | 0.0156 | 0.49 |
| Guanidoacetic acid | 0.0164 | 0.23 |
| Succinic acid | 0.0164 | 0.36 |
| Ascorbic acid | 0.0199 | 0.59 |
| Cystathionine | 0.0223 | -1.00 |
| Dihydrouracil | 0.0244 | -0.63 |
| 4-Methyl-2-oxopentanoic acid | 0.0255 | 0.36 |
| N-Methyl-glutamic acid | 0.0259 | -0.27 |
| Pipecolic acid | 0.0322 | 0.46 |
| Glyceric acid | 0.0334 | 0.21 |
| CAR(14:0) | 0.0361 | 0.59 |
| Lysine | 0.0361 | -0.28 |
| NADPH | 0.0409 | 0.58 |
| Citrulline | 0.0414 | -0.19 |
| Cytosine | 0.0443 | -0.34 |
| Lactic acid | 0.0472 | 0.21 |
| **Choroid** |  |  |

| Metabolite | P-value | Log2(FoldChange) |
| --- | --- | --- |
| CAR(8:0) | 5.58E-06 * | -2.37 |
| 5-Methylthioadenosine | 4.41E-05 * | 1.43 |
| Deoxyuridine | 9.38E-05 * | 2.48 |
| CAR(6:0) | 0.0001 * | -1.56 |
| Uric acid | 0.0002 * | 1.42 |
| CAR(10:0) | 0.0002 * | -1.87 |
| CAR(12:0) | 0.0004 * | -1.12 |
| S-Adenosylhomocysteine | 0.0005 * | 1.05 |
| 3-Methylhistamine | 0.0006 * | 1.96 |
| CAR(5:0) | 0.0007 * | -1.55 |
| Riboflavin | 0.0026 * | 0.84 |
| N-Acetylserotonin | 0.0049 | -0.67 |
| N-Acetylcysteine | 0.0051 | 0.99 |
| CAR(4:0) | 0.0071 | -0.85 |
| CAR(3:0) | 0.0088 | -0.89 |
| cAMP | 0.0089 | 1.19 |
| Tyrosine | 0.0099 | 0.50 |
| Guanine | 0.0105 | -0.85 |
| Indole-3-lactic acid | 0.0136 | 1.69 |
| Glyceraldehyde 3-phosphate | 0.0146 | -1.38 |
| CAR(20:4) | 0.0166 | -0.90 |
| S-Adenosylmethionine | 0.0179 | 0.85 |
| 4-Methyl-2-oxopentanoic acid | 0.0191 | 0.40 |
| Allantoin | 0.0210 | 1.13 |
| Acetyl-L-carnitine | 0.0223 | -0.56 |
| Cholic acid | 0.0228 | 1.29 |
| Adenine deoxyribonucleoside | 0.0234 | -0.90 |
| Pyruvic acid | 0.0239 | -0.52 |
| Melatonin | 0.0243 | -1.41 |
| Tryptophan | 0.0249 | -0.92 |
| CDP-Ribitol | 0.0311 | -0.69 |
| CAR(18:1) | 0.0349 | 1.02 |
| dAMP | 0.0357 | -0.87 |
| Guanosine | 0.0375 | -0.86 |
| Oxalic acid | 0.0414 | -0.55 |
| Gamma-glutamylcysteine | 0.0453 | 0.65 |
| UDP-glucuronic acid | 0.0497 | -1.74 |

**Table 4.** Metabolite differences in human RPE/choroid during aging and AMD. Data corresponds to Figure 3. Asterisks (*) indicate significance according to Benjamini-Hochberg FDR correction.

| Human Extramacular Choroid: Aging and AMD |  |  |
| --- | --- | --- |
| Young vs Aged |  |  |
| Metabolite | P-value | Log2(FoldChange) |
| Uric acid | 6.58E-07 * | 1.19 |
| TMAO | 4.96E-05 * | 4.14 |
| Quinic acid | 0.0001 * | 3.47 |
| Guanidinosuccinic acid | 0.0003 | 3.54 |
| Gluconic acid | 0.0008 | 1.54 |
| Pyruvic acid | 0.0017 | 1.22 |
| 4-Methyl-2-oxopentanoic acid | 0.0019 | 0.86 |
| Tryptophan | 0.0021 | 0.66 |
| N-Methyl-glutamic acid | 0.0025 | 0.63 |
| N-Lactoyl-Phenylalanine | 0.0030 | 0.69 |
| Indoxyl sulfate | 0.0030 | 2.01 |
| 3-Methyl-2-oxobutanoic acid | 0.0046 | 0.85 |
| Dimethylarginine | 0.0046 | 0.89 |
| Adenosine | 0.0048 | 0.85 |
| Glycerol 3-phosphate | 0.0059 | -0.70 |
| sn-Glycerol 3-phosphate | 0.0060 | -0.70 |
| Glucose | 0.0060 | 1.95 |
| Cholic acid | 0.0061 | 2.66 |
| 3-Hydroxypropionic acid | 0.0067 | 0.53 |
| Lactic acid | 0.0067 | 0.53 |
| 2-Hydroxybutyric acid | 0.0086 | 0.85 |
| Creatinine | 0.0099 | 0.95 |
| Cortisol | 0.0109 | 1.48 |
| Allantoin | 0.0135 | 1.10 |
| Phenylalanine | 0.0229 | 0.39 |
| S-Adenosylmethionine | 0.0281 | 0.41 |
| Heptadecanoic acid | 0.0350 | 0.43 |
| N-Acetylalanine | 0.0360 | 0.38 |
| Serine | 0.0432 | 0.33 |
| Vanillylmandelic acid | 0.0465 | 0.95 |
| N-Formylmethionine | 0.0490 | 0.42 |
| NADP+ | 0.0500 | -0.61 |
| **Aged vs Early/Intermediate AMD** |  |  |
| **Metabolite** | **P-value** | **Log2(FoldChange)** |
| N-Acetylneuraminic acid | 0.0260 | 0.51 |
| Glutamic acid | 0.0282 | 0.35 |
| Histidine | 0.0311 | -0.22 |
| 3-Methylhistamine | 0.0425 | -0.38 |
| N-Formylmethionine | 0.0476 | -0.27 |
| CAR(20:4) | 0.0491 | 0.59 |
| **Aged vs Geographic Atrophy** |  |  |
| **Metabolite** | **P-value** | **Log2(FoldChange)** |
| Azelaic acid | 0.0060 | -0.54 |
| Nonanoic acid | 0.0145 | -0.68 |
| Heneicosanoic acid | 0.0199 | -0.59 |
| Stearic acid | 0.0199 | -0.59 |
| Heptadecanoic acid | 0.0323 | -0.46 |
| Arachidic acid | 0.0359 | -0.61 |
| N-Acetylglutamic acid | 0.0364 | 0.40 |
| Glycine | 0.0413 | 0.43 |
| S-Adenosylhomocysteine | 0.0444 | 0.48 |
| CAR(5:0-DC) | 0.0484 | -1.17 |
| Pentadecanoic acid | 0.0491 | -0.34 |
| **Aged vs Macular Neovascularization** |  |  |
| **Metabolite** | **P-value** | **Log2(FoldChange)** |
| dAMP | 0.0018 | -0.76 |
| Aminolevulinic acid | 0.0032 | 0.20 |
| TMP | 0.0059 | -1.14 |
| Glutamine | 0.0129 | 0.25 |
| GMP | 0.0157 | -0.68 |
| CAR(20:4) | 0.0213 | -0.79 |
| AMP | 0.0244 | -0.61 |
| Adenine deoxyribonucleoside | 0.0286 | -0.51 |
| GDP | 0.0350 | -0.37 |
| CDP | 0.0395 | -0.80 |

### Human: TMAO and Uric Acid

Two particularly interesting metabolites emerged from the results of the LC-MS analysis: trimethylamine N-oxide (TMAO) and uric acid (UA). TMAO is a small organic compound derived from microbiome-dependent metabolism of carnitines and choline, while UA is the end product of purine catabolism. To further examine TMAO and UA in the context of the choroid, functional studies were undertaken using cell culture monolayers. Each metabolite was also measured again using absolute quantification in retina, RPE/choroid, and serum from 4 donors with high TMAO and high UA and 4 with low TMAO and low UA, which confirmed the high and low categorizations found in the original relative LC-MS data (Sup. Table 2).

To determine if either TMAO or UA altered cell migration, wound healing assays were conducted using C166 endothelial cells as described in the methods. Succinylated concanavalin A (sConA) was used as a negative control, which slows migration and aggregates C166 cells over time (unpublished). This is shown by the negative percentage closure in the sConA-treated cells (Fig. 4B). When compared to respective controls, neither concentration of TMAO displayed a significant difference in closure speed at any timepoint. Cells treated with 100 µg/mL uric acid also did not show altered migration. However, cells treated with 200 µg/mL uric acid closed significantly slower than the NaOH vehicle control (Fig. 4A), with a statistically significant difference at every timepoint after 0h. Representative wound area analysis images are shown in Figure 4B, demonstrating a similarly sized initial wound area and slower closure of the 200 µg/mL uric acid treated cells.

**Figure 4.**
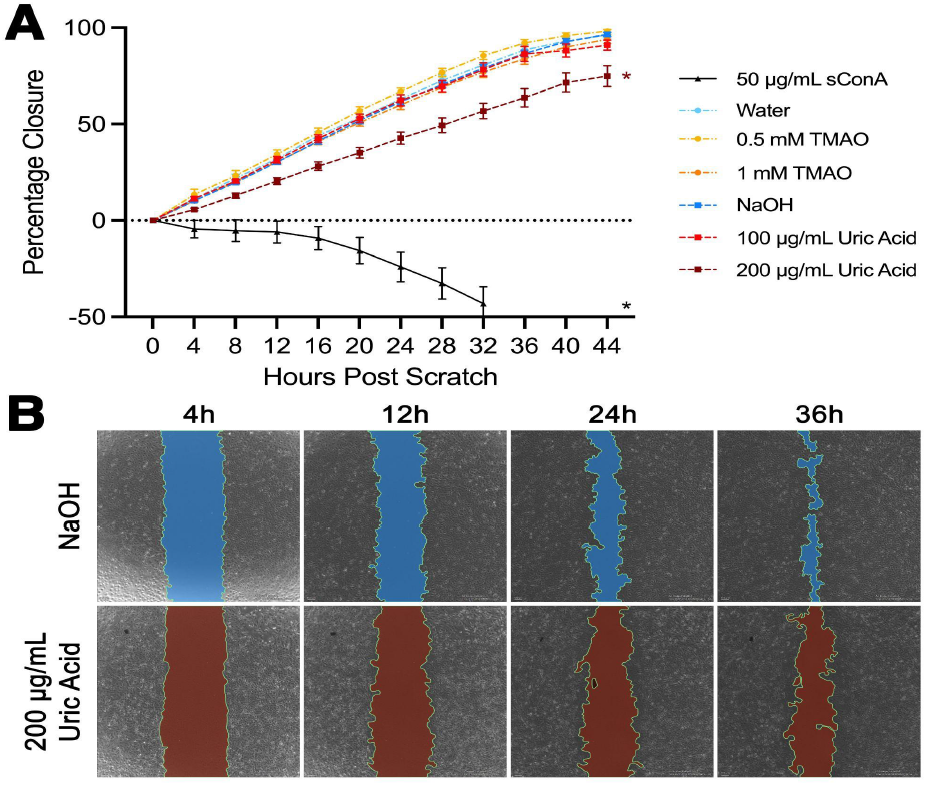
Wound closure in C166 cells treated with TMAO and Uric Acid. Panel A displays the percentage wound closure over 44 hours. For statistical analysis, each TMAO or uric acid treatment was compared to the appropriate vehicle control (water and NaOH, respectively). Representative images of NaOH vehicle control and 200 µg/mL uric acid treated cells at 4, 12, 24, and 36 hours are shown in (B), with wound area shown in blue or red for NaOH or 200 µg/mL uric acid respectively. Asterisks indicate a p-value of 0.05 or less. Percentage closure of sConA and 200 µg/mL uric acid treated cells was significantly different from controls at every timepoint. Each point represents the mean of 9 replicates, while error bars represent the standard error of the mean.

## DISCUSSION

In this study, we sought to profile the metabolome of the peripheral RPE and choroid from a large set of well characterized donor eyes. Because extramacular tissue was available for these experiments, we performed initial comparison of macula and extramacular RPE/choroid from a separate cohort of 10 donors. This experiment showed relatively minimal differences between the two, with only 11 significantly different metabolites and no pathways reaching significance. This result showed that extramacular RPE/choroid could be used to examine metabolic changes in the eye in aging and disease.

Metabolomic profiling was performed on a set of 87 RPE/choroid punches of different ages and disease states. It is notable that the most significant differences were observed between young and healthy aged controls, not between healthy controls and any stage of AMD. This is reflected in both the number of significantly different metabolites and pathways, the magnitude of fold changes and pathway impacts, and the degree of significance. There is also overlap in divergent metabolites and pathways between the human aging and mice aging comparison. Overlapping significant pathways include glycerophospholipid metabolism, glycerolipid metabolism, purine metabolism, sphingolipid metabolism, arginine and proline metabolism, histidine metabolism, and nicotinate and nicotinamide metabolism. There are an additional 8 pathways in humans and 21 in mice that are significant only in one of the tested species. This could be due to inherent differences in metabolism between humans and mice, or environmental variation such as diet. An example of a difference likely attributable to diet is the presence of significantly increased TMAO in humans, but not in mice, which is discussed in further detail below.

Though relatively modest, some metabolic changes were observed between healthy aged human choroids and various stages of AMD. Early/intermediate AMD choroids have significantly different nitrogen metabolism, porphyrin metabolism, and glycine/serine/threonine metabolism, though only the latter two have a Pathway Impact score of above 0. Porphyrin metabolism was also significantly different in MNV compared to aged controls, along with steroid hormone biosynthesis and pyrimidine metabolism. No pathways were different between GA and aged controls. The results of this analysis show that there are metabolic changes in the choroid of eyes with varying stages of AMD, even in the extramacular regions that are spatially removed from the primary site of pathogenesis. The finding of increased N-Acetylneuraminic acid (NeuNAc) in early AMD compared to controls was particularly interesting in light of NeuNAc-containing glycans in AMD-associated deposits and its role in binding Factor H (32,33).

Two metabolites of particular interest emerged from human young vs aged choroid comparison: trimethylamine N-oxide (TMAO) and uric acid. In humans, the large majority of TMAO is derived from the microbiome (34). L-carnitine and choline from the diet are catabolized to TMA in microbiota. TMA is then released in the colon and can either be degraded to other compounds such as methylamine, dimethylamine, and ammonia or absorbed by the bloodstream and oxidized into TMAO by flavin monooxygenases. TMAO can then be accumulated in tissues as an osmolyte or cleared by the kidneys and excreted in the urine. High plasma concentration of TMAO has been associated with an increased risk of cardiovascular disease and cardiac events (34–41). TMAO has also been demonstrated to promote endothelial dysfunction through increased oxidative stress in murine and human endothelial cells (42–44). In the current study, TMAO was elevated more than 17-fold in aged human donors, with a p-value of 4.95E-05. This was the largest fold change and the second smallest p-value of all metabolites in this comparison. TMAO was not found to be significantly different between young and aged mice. This could be due to many factors, such as an identical diet between all animals, older animals not being old enough to accumulate TMAO, different flora in mouse and human, or the lack of diet-based sources of TMAO precursors. Additionally, we evaluated whether TMAO levels differed between aged donors (unaffected and affected) with and without a determination of sepsis. No significant difference in the abundance of this metabolite was observed in relation to this diagnosis. While initial functional experiments with TMAO-treated C166 cells did not display a change in migration, further studies of the impact on this metabolite and its precursors on human endothelial cells is warranted.

Uric acid is the terminal product of purine catabolism. Guanosine and adenosine monophosphates are degraded to xanthine, which is then oxygenated by xanthine oxidase to produce uric acid. Uric acid has low solubility in water and generally exists in the body as urate (ionized salt form). Uric acid is excreted in urine via the kidneys. At physiological levels, uric acid is a potent antioxidant that can act as a reducing agent to neutralize reactive oxygen species. High blood concentrations of uric acid (hyperuricemia) can cause gout and nephrolithiasis (45), and are associated with a variety of other conditions such as type 2 diabetes, fatty liver, chronic kidney disease, and cardiovascular disease (46–52). Uric acid has also been associated with accelerated aging (53) and higher all-cause mortality (54). Elevated uric acid has been demonstrated to cause endothelial cell dysfunction through multiple mechanisms. Endothelial dysfunction can be induced by reduced nitric oxide synthesis, which hyperuricemia can cause by impairing eNOS activity, reducing the supply of L-arginine, and increasing superoxide production (55). This increased oxidative stress can also cause endothelial insulin resistance, endothelial to mesenchymal transition, and injury via inflammasome activation.

In aged human choroids, uric acid was elevated 2.28-fold with a p-value of 6.58E-07. In aged mouse choroids, uric acid was elevated 2.68-fold (p=2.05E-04), suggesting that this finding is generalizable. C166 cell monolayers treated with uric acid exhibited impaired migration compared to vehicle controls.

This study has advantages and limitations. From the mouse postmortem metabolome profiling, it is clear that there are a very large number of changes in small molecule abundance after death. The timepoint of 7 hours postmortem is close to the maximum death to preservation time in the human samples used, so this experiment provides an overview of which metabolites and pathways are least stable in the RPE/choroid over time. Though there is significant change in the metabolome during the postmortem time, each group of human RPE/choroids that were compared had similar average death to preservation times with a presumably similar impact on metabolite degradation. Levels of TMAO and uric acid were not related to postmortem interval in human choroid; moreover, uric acid was also elevated in aged mouse choroids collected immediately after death. Therefore, although the data herein is not reflective of a living eye, each group can still be compared to provide initial insights into human RPE/choroid metabolomes with what is currently the lowest postmortem times achievable. However, this is an inherent limitation to studying human tissue. In addition, extramacular tissue available for this experiment was a surrogate for macular biochemistry. While the RPE/choroid from the extramacular region was similar to that of the macula, in the context of atrophy or other pathology restricted to the macula, studies that include the macular lesions would be beneficial.

In conclusion, this study provides initial insight into the metabolic state of postmortem human RPE/choroid tissue at varying ages and states of macular degeneration. Differences between young and aged RPE/choroid were notable, and select metabolites (TMAO and uric acid) were functionally examined, with uric acid showing an impact on endothelial cell behavior. This work provides a starting point for understanding both challenges and new insights from examining metabolism in human postmortem ocular tissues. We are hopeful that providing the raw data from these experiments to the community will accelerate understanding and progress in promoting choroidal health in aging and AMD.

## Supporting information

Navratil choroidal metabolomics-supplemental

## ACKNOWLEDGEMENTS

Funded in part by NIH Grants EY-033308, EY-024605, EY-025580, and DK138664. LC-MS Metabolomics analysis and initial data processing was conducted by the University of Iowa Fraternal Order of Eagles Diabetes Research Center Metabolomics core. The authors thank the Iowa Lions Eye Bank, the eye donors, and their families for their invaluable contribution to this research.

## Notes

### Competing Interest Statement

The authors have declared no competing interest.

