## Supplementary material for "Metabolic Analysis of Human Retinal Pigment Epithelium and Choroid Tissue in Aging and Macular Degeneration": Navratil choroidal metabolomics-supplemental

**Supplementary Table 1.** Human Donor Demographic Characteristics. Includes age, sex, ocular disease category, death to preservation time (rounded to the nearest half hour), and cause of death.

| Donor # | Age | Sex | Disease State | Death to Preservation Time (hours:minutes) | Cause of Death |
|---|---|---|---|---|---|
| Macula vs Periphery Cohort | | | | | |
| 1 | 13 | Male | Control | 6:30 | Cardiopulmonary Failure, Lymphoblastic Leukemia |
| 2 | 15 | Female | Control | 5:30 | Mesenchymal Chondrosarcoma with Mets |
| 3 | 49 | Female | Control | 6:00 | Adenocarcinoma with Mets to Bone and Sepsis |
| 4 | 54 | Male | Control | 7:00 | Pancreatic cancer |
| 5 | 64 | Female | Control | 5:00 | Lung Cancer with Mets |
| 6 | 66 | Male | Control | 7:30 | Acute Renal Failure and Sepsis |
| 7 | 68 | Female | Control | 4:00 | Sepsis |
| 8 | 72 | Male | Control | 7:00 | Septic Shock |
| 9 | 73 | Female | Control | 7:30 | Cardiogenic Shock |
| 10 | 77 | Male | Control | 5:30 | Respiratory Arrest |
| Aging and AMD Cohort | | | | | |
| 11 | 21 | Female | Young | 5:30 | Hodgkin's Lymphoma |

| | | | | | |
|---|---|---|---|---|---|
| 12 | 22 | Female | Young | 7:00 | Cardiopulmonary Arrest/Pneumonia |
| 13 | 23 | Male | Young | 6:00 | Respiratory Failure/Testicular Cancer |
| 14 | 32 | Male | Young | 6:30 | Leukemia/Cardiac Arrest |
| 15 | 36 | Female | Young | 6:00 | Drug Overdose |
| 16 | 40 | Male | Young | 6:00 | Alcoholic Liver Failure/Septic Shock |
| 17 | 42 | Male | Young | 6:00 | Sepsis |
| 18 | 42 | Female | Young | 6:30 | Acute Liver and Renal Failure |
| 19 | 70 | Male | Control | 6:00 | Unavailable |
| 20 | 70 | Male | Control | 6:00 | Intrahepatic Bile Duct Carcinoma with Mets |
| 21 | 70 | Male | Control | 5:30 | Acute Hypoxic Respiratory Failure |
| 22 | 71 | Female | Control | 6:30 | Liver Cirrhosis/Sepsis |
| 23 | 72 | Female | Control | 6:30 | Sepsis |
| 24 | 76 | Female | Control | 5:00 | Hyperkalemia/Cardiac Arrest |
| 25 | 76 | Male | Control | 6:00 | Hypoxic Respiratory Failure Secondary to Intracerebral Hemorrhage |
| 26 | 77 | Male | Control | 5:30 | Renal Failure |
| 27 | 77 | Female | Control | 6:30 | Cardiogenic Shock/Sepsis |
| 28 | 78 | Male | Control | 7:30 | Heart Failure & Pneumonia |

| | | | | | |
|---|---|---|---|---|---|
| 29 | 79 | Male | Control | 6:30 | Sepsis |
| 30 | 79 | Male | Control | 5:00 | Cardiopulmonary Arrest |
| 31 | 81 | Female | Control | 5:00 | Sepsis |
| 32 | 82 | Female | Control | 5:30 | Dry Gangrene/Supratherapeutic INR & Upper GI Bleed |
| 33 | 83 | Female | Control | 6:30 | Gastric Bowel Perforation |
| 34 | 83 | Male | Control | 6:00 | Cardiopulmonary Arrest |
| 35 | 84 | Female | Control | 5:30 | Heart Attack |
| 36 | 84 | Male | Control | 4:00 | Prostate Cancer with Metastasis |
| 37 | 85 | Male | Control | 6:30 | Cardiogenic Shock |
| 38 | 86 | Female | Control | 5:00 | Respiratory Failure |
| 39 | 86 | Male | Control | 5:00 | Aortic Stenosis |
| 40 | 87 | Female | Control | 5:30 | Stroke |
| 41 | 87 | Female | Control | 5:30 | Stroke |
| 42 | 87 | Female | Control | 7:00 | Septic Shock |
| 43 | 88 | Male | Control | 3:00 | Unavailable |
| 44 | 89 | Female | Control | 5:00 | Acute Subdural Hematoma |
| 45 | 89 | Female | Control | 3:30 | Pneumonia |
| 46 | 89 | Male | Control | 6:00 | End Stage Renal Disease |
| 47 | 90 | Male | Control | 6:00 | Cancer |

| | | | | | |
|---|---|---|---|---|---|
| 48 | 90 | Female | Control | 6:00 | Cholecystitis |
| 49 | 91 | Female | Control | 8:00 | Malnutrition |
| 50 | 91 | Female | Control | 6:30 | Pulmonary Embolism |
| 51 | 92 | Male | Control | 6:00 | Natural Causes |
| 52 | 92 | Female | Control | 5:00 | Respiratory Failure Secondary to Ischemic Stroke |
| 53 | 94 | Male | Control | 7:00 | Vagal Episode |
| 54 | 96 | Female | Control | 5:30 | COPD |
| 55 | 97 | Female | Control | 7:00 | Sepsis/Right Femur Fracture |
| 56 | 71 | Male | AMD | 4:30 | Cardiac Arrest |
| 57 | 73 | Male | AMD | 6:30 | Respiratory Failure with Hypoxia |
| 58 | 76 | Female | AMD | 8:00 | Unavailable |
| 59 | 77 | Male | AMD | 6:30 | Pneumonia |
| 60 | 79 | Male | AMD | 7:00 | Pancreatic Cancer |
| 61 | 79 | Female | AMD | 6:30 | Congestive Heart Failure |
| 62 | 83 | Male | AMD | 5:30 | Motor Vehicle Collision |
| 63 | 83 | Male | AMD | 7:00 | Motor Vehicle Accident |
| 64 | 84 | Male | AMD | 6:00 | Stage 4 Prostate Cancer |
| 65 | 85 | Male | AMD | 6:00 | Acute Coronary Syndrome |
| 66 | 85 | Female | AMD | 4:30 | Cardiogenic Shock |

| | | | | | |
|---|---|---|---|---|---|
| 67 | 86 | Male | AMD | 7:30 | COPD |
| 68 | 89 | Female | AMD | 4:30 | Cerebrovascular Accident |
| 69 | 89 | Female | AMD | 4:00 | Unavailable |
| 70 | 89 | Male | AMD | 6:30 | Congestive Heart Failure |
| 71 | 89 | Female | AMD | 6:30 | Pneumonia |
| 72 | 89 | Female | AMD | 7:30 | Bladder Cancer with Bone Mets |
| 73 | 92 | Female | AMD | 5:30 | Lung Cancer |
| 74 | 92 | Male | AMD | 4:30 | Multi-Organ Failure, Respiratory Failure |
| 75 | 93 | Female | AMD | 4:30 | Cardiac Arrest |
| 76 | 96 | Female | AMD | 5:30 | Sepsis |
| 77 | 76 | Female | GA | 4:00 | Unavailable |
| 78 | 87 | Female | GA | 5:00 | Leukemia, Acute Renal Failure |
| 79 | 87 | Female | GA | 5:00 | Cardiac Arrest |
| 80 | 87 | Female | GA | 4:30 | Renal Failure |
| 81 | 92 | Female | GA | 5:30 | Respiratory Distress |
| 82 a | 93 | Female | GA | 8:00 | Cardiogenic Shock |
| 82 b | | | MNV | 7:30 | |
| 83 | 97 | Female | GA | 6:00 | Cerebral Vascular Accident |

| | | | | | |
|---|---|---|---|---|---|
| 84 | 70 | Female | MNV | 8:00 | Cardiac Arrest |
| 85 | 72 | Female | MNV | 6:00 | Post Surgical Complications - Bowel Resection |
| 86 | 77 | Male | MNV | 6:00 | Unavailable |
| 87 | 80 | Male | MNV | 4:30 | Respiratory Arrest |
| 88 | 81 | Male | MNV | 7:30 | Cardiac Arrest |
| 89 | 82 | Female | MNV | 6:00 | Acute Renal Failure |
| 90 | 83 | Female | MNV | 4:30 | Subdural Hematoma |
| 91 | 83 | Female | MNV | 5:00 | Metastatic Melanoma |
| 92 | 85 | Female | MNV | 5:00 | Aortic Aneurysm |
| 93 | 87 | Female | MNV | 4:00 | Small Bowel Obstruction |
| 94 | 92 | Male | MNV | 5:00 | Sepsis - Pneumonia |
| 95 | 92 | Female | MNV | 6:30 | Sepsis |
| 96 | 97 | Male | MNV | 4:00 | Cardiac Arrest |

**Supplemental Table 2.** Absolute TMAO and Uric Acid Quantification. Categorization of high or low is based on results from the initial relative LC-MS metabolomics data.

| TMAO Absolute Quantity | | | | |
|---|---|---|---|---|
| Category (High or Low TMAO) | Donor ID | RPE/Choroid (µg/mg sample) | Retina (µg/mg sample) | Serum (µg/mL sample) |
| Low | 11 | 0.016 | 0.024 | 12.941 |
| | 13 | 0.015 | 0.031 | 3.387 |
| | 16 | 0.025 | 0.028 | 10.887 |
| | 18 | 0.019 | 0.048 | 13.643 |
| High | 31 | 0.285 | 0.502 | 636.552 |
| | 35 | 0.109 | 1.406 | 127.489 |
| | 36 | 0.119 | 0.244 | 191.676 |
| | 51 | 0.307 | 0.340 | 622.375 |
| Uric Acid Absolute Quantity | | | | |
| Category (High or Low Uric Acid) | Donor ID | RPE/Choroid (µg/mg sample) | Retina (µg/mg sample) | Serum (µg/mL sample) |
| Low | 11 | 1.16 | 25.70 | 34053.87 |
| | 13 | 3.18 | 7.750 | 28426.45 |
| | 16 | 5.51 | 5.74 | 17807.74 |
| | 18 | 10.13 | 42.09 | 69116.16 |
| High | 31 | 15.14 | 14.81 | 121632.61 |
| | 35 | 42.97 | 329.69 | 116308.74 |
| | 36 | 43.83 | 86.52 | 143689.30 |
| | 51 | 25.03 | 22.53 | 110119.52 |

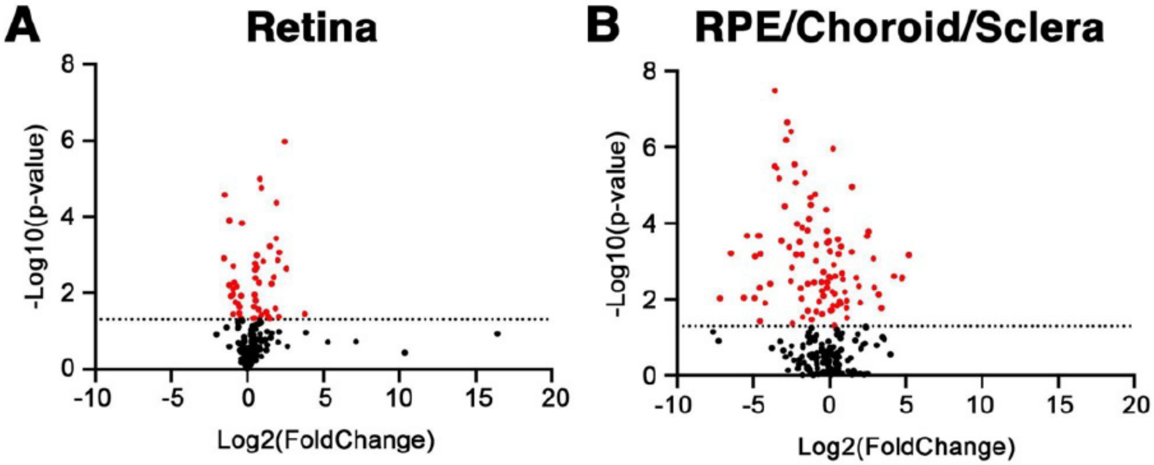


**Supplementary Figure 1**. Metabolic changes in the mouse RPE/choroid and neural retina at 7 hours postmortem. A volcano plot representing metabolite changes in the retina after 7 hours is shown in A, while the same analysis for RPE/choroid/sclera is shown in B. Dotted lines indicate a p-value of 0.05.
